# Individual differences in locomotor function predict the capacity to reduce asymmetry and modify the energetic cost of walking post-stroke

**DOI:** 10.1101/197582

**Authors:** Natalia Sánchez, James M. Finley

**Affiliations:** Division of Biokinesiology and Physical Therapy, University of Southern California, Los Angeles, CA, 90033; Department of Biomedical Engineering, University of Southern California, Los Angeles, CA, 90089; Neuroscience Graduate Program, University of Southern California, Los Angeles, CA, 90089

## Abstract

Changes in the control of the lower extremities post-stroke lead to persistent biomechanical asymmetries during walking. These asymmetries are associated with an increase in energetic cost, leading to the possibility that reduction of asymmetry can improve economy. However, the influence of asymmetry on economy may depend on the direction and cause of asymmetry. For example, impairments with paretic limb advancement may result in shorter paretic steps while deficits in paretic support or propulsion result in shorter non-paretic steps. Given differences in the underlying impairments responsible for each type of step length asymmetry, the capacity to reduce asymmetry, and the associated changes in energetic cost may not be consistent across this population. Here, we identified factors explaining individual differences in the capacity to voluntarily reduce step length asymmetry and modify energetic cost during walking. Twenty-four individuals post-stroke walked on a treadmill with visual feedback of their step lengths to aid explicit modification of asymmetry. We found that individuals who naturally took longer paretic steps had a greater capacity to reduce asymmetry, and were better able to transfer the effects of training to over-ground walking. In addition, baseline energetic cost was negatively correlated with reductions in cost, such that participants with a more economical gait were more likely to reduce energetic cost by improving symmetry. These results demonstrate that many stroke survivors retain the capacity to voluntarily walk more symmetrically on a treadmill and over-ground. However, whether reductions in asymmetry reduce metabolic cost depends on individual differences in impairments affecting locomotor function.

## Introduction

One of the major goals of post-stroke rehabilitation is to improve walking ability^1,2^. Post-stroke changes in lower extremity function lead to persistent biomechanical abnormalities during walking including but not limited to decreased paretic stance time^3^, and asymmetric step lengths^4–7^. Previous research has shown positive associations between spatiotemporal asymmetries and the metabolic cost of walking^8–10^, and in people post-stroke, this increased cost may be due in part to sub-optimal coordination of leading limb contact and trailing limb push-off forces^10,11^. Therefore, it is possible that reducing asymmetry may reduce the metabolic burden of walking post-stroke. Alternatively, people post-stroke may naturally adopt asymmetric walking patterns because this strategy is energetically optimal given their level of impairment. However, these possibilities have yet to be systematically investigated.

Due to heterogeneity in the types of motor impairments that occur post-stroke, the direction and magnitude of step length asymmetry are not uniform across the entire post-stroke population. Whereas most individuals shorten stance time and increase swing time on the paretic side^3,12,13^, the direction of the spatial asymmetry is highly heterogeneous^5,14,15^. Reductions in paretic propulsion^5,14^ and hip extension^16^ manifest as shorter non-paretic steps while dominant impairments with paretic limb advancement^5,17^ may result in shorter paretic steps. Given the differences in the mechanisms underlying each type of asymmetry, the capacity to reduce asymmetry may not be consistent across the entire post-stroke population. Moreover, these differential impairments may influence how metabolic cost changes in response to reductions in asymmetry: individuals with shorter non-paretic steps may only need to increase paretic stance duration^18^ whereas individuals with shorter paretic steps may rely on strategies such as hip hiking and circumduction to advance the paretic limb, which could increase metabolic cost^19–23^.

Here, we sought to determine: 1) the factors that predict whether people post-stroke retain the capacity to reduce step length asymmetry, 2 voluntarily) how changes in asymmetry impact metabolic cost and 3) whether voluntary attempts to reduce asymmetry during treadmill walking transfer to over-ground walking. We addressed these aims through the use of a single session, biofeedback-based, treadmill training paradigm which enabled participants to reduce step length asymmetry explicitly. We hypothesized that the baseline direction of asymmetry will influence the capacity to reduce asymmetry voluntarily and that reductions in asymmetry would be correlated with reductions metabolic cost. We also hypothesized that the direction of the baseline asymmetry would influence how well reductions in asymmetry transfer to over-ground walking. Ultimately, a better understanding of how motor impairments influence the ability to reduce asymmetry can facilitate the design of personalized interventions to improve locomotor function post-stroke.

## Methods

Twenty-four chronic stroke survivors (Table 1) participated in this study. Study inclusion criteria were: 1) chronic hemiparesis (time since stroke > 6 months) due to a single stroke, 2) ability to walk on the treadmill continuously for five minutes, 3) ability to walk over-ground independently or with use of a cane, 4) no concurrent neurological disorders or orthopedic conditions that interfered with their ability to walk, and 5) the ability to provide informed consent. Consistent with previous studies^24,25^, participants were instructed to lightly touch a handrail placed in front of them to aid balance and prevent drift on the treadmill. All procedures conformed to the principles set forth in the Declaration of Helsinki and were approved by the University of Southern California’s Institutional Review Board.

**Table 1:** Participant demographics

| ID | Sex | Age | Months Since<br>Stroke | Paretic<br>Side | FM | OG<br>Speed (m/s) | OG Asym<br>(mm) | TM<br>Speed<br>(m/s) | TM Asym<br>(mm) | Lesion<br>Type |
| --- | --- | --- | --- | --- | --- | --- | --- | --- | --- | --- |
| 1<sup>b</sup> | M | 58 | 83 | Left | 20 | 0.76 (10mwt) | -26.12 | 0.48 | **6.50** | Hem |
| 2<sup>b</sup> | M | 49 | 43 | Right | 25 | 0.95 (10mwt) | 9.13 | 0.55 | 6.87 | Isch |
| 3 | M | 73 | 143 | Left | 25 | 0.8 (10mwt) | 52.40 | 0.6 | 8.62 | Isch |
| 4 | F | 58 | 44 | Right | 23 | 1.01 | -24.23 | 0.65 | -74.87 | Isch |
| 5 | M | 45 | 53 | Left | 25 | 0.93 | 21.96 | 0.7 | 11.32 | Isch |
| 6<sup>c,b</sup> | F | 64 | 369 | Right | 18 | 0.82 | -118.55 | 0.6 | **76.87** | Isch |
| 7 | M | 69 | 89 | Left | 23 | 0.37 | 209.50 | 0.37 | 117.18 | Isch |
| 8 | M | 76 | 73 | Right | 27 | 0.95 | -76.97 | 0.7 | **10.35** | Isch |
| 9 | M | 54 | 41 | Right | 24 | 0.78 | -62.21 | 0.65 | -30.72 | Hem |
| 10<sup>b</sup> | F | 39 | 324 | Right | 24 | 0.82 (10mwt) | -72.96 | 0.5 | **40.60** | Isch |
| 11 | M | 58 | 43 | Right | 23 | 0.65 | 36.08 | 0.4 | 21.60 | Hem |
| 12 | F | 56 | 85 | Right | 27 | 0.58 | -158.07 | 0.58 | -148.27 | Hem |
| 13 | M | 63 | 52 | Left | 23 | 0.92 | -3.70 | 0.65 | -55.07 | Isch |
| 14<sup>c,b</sup> | F | 72 | 154 | Right | 10 | 0.49 | 126.91 | 0.32 | **-40.54** | Hem |
| 15 | M | 67 | 116 | Right | 26 | 0.98 | -50.49 | 0.8 | -18.25 | Hem |
| 16<sup>c</sup> | M | 45 | 118 | Left | 7 | 0.14 | 29.71 | 0.33 | 198.39 | Isch |
| 17 | M | 61 | 34 | Left | 16 | 0.27 | 13.61 | 0.25 | **-69.18** | Isch |
| 18 | M | 60 | 169 | Right | 22 | 0.6 | -72.70 | 0.45 | -61.78 | Hem |
| 19 | M | 33 | 37 | Right | 28 | 0.95 | 14.73 | 0.95 | 44.87 | Isch |
| 20 | M | 61 | 52 | Right | 32 | 1 | -49.09 | 0.85 | -84.73 | Isch |
| 21<sup>b</sup> | M | 56 | 88 | Right | 19 | 0.42 | -12.32 | 0.38 | **68.26** | Hem |
| 22<sup>c</sup> | M | 62 | 96 | Left | 18 | 0.48 | -41.93 | 0.35 | **23.68** | Hem |
| 23<sup>b</sup> | F | 28 | 25 | Left | 19 | 0.17 | 8.81 | 0.13 | 35.98 | Hem |
| 24 | M | 55 | 15 | Left | 27 | 0.85 | -90.45 | 0.65 | -166.66 | Isch |
| **Average** | **NA** | **56.75** | **97.75** | **NA** | **22.13** | **0.67** |  | **0.53** |  | **NA** |

<sup>b</sup> Participant wore an ankle brace during the experiment. <sup>c</sup> Participant used a cane during over-ground walking. Isch: Ischemic stroke. Hem: Hemorrhagic stroke. For four participants, gait speed was assessed using the 10-meter walk test (10mwt) instead of the 6-minute walk test. Bold asymmetries indicate those participants whose asymmetry direction was different over-ground and on the treadmill. It should be noted that most of these individuals used an assistive device when walking over-ground.

### Experimental Protocol

The full protocol consisted of a set of clinical assessments and a combination of trials over-ground and on the treadmill (Figure 1). The lower extremity portion of the Fugl-Meyer Assessment^26^ was performed to assess lower extremity motor impairment. Initial evaluation of walking function was done using the 6-minute walk test. The experiment began with 4-6 passes of over-ground walking over a 10m walkway at each participant’s self-selected speed (*OG BASE*), and this was followed by a treadmill familiarization trial. During all treadmill trials, participants wore a harness to prevent falls, without providing any body weight support. During this trial, the speed of the treadmill was gradually adjusted using the staircase method^27^ until participants achieved their comfortable walking speed. In the final minute of this five-minute trial, we introduced the visual feedback and participants practiced matching step length targets for one minute. They subsequently completed a 5-minute walking trial on the treadmill (*BASELINE*) where we measured their baseline metabolic cost and step lengths. After *BASELINE*, participants walked for five minutes with feedback of their natural step lengths (*BASELINE+FBK*) to measure any potential changes in spatiotemporal control or metabolic cost due to the accurate foot placement requirements. For the subsequent *SYMMETRY* trial, participants were instructed to walk with equal step lengths, by lengthening their shorter step to match their longer step. Lastly, participants transitioned to walking over-ground to assess transfer (*OG POST*). Sitting breaks of at least five minutes were provided between all trials, and blood pressure and heart rate were measured for safety and to ensure that resting conditions were achieved before beginning the next trial.

**Figure 1.**
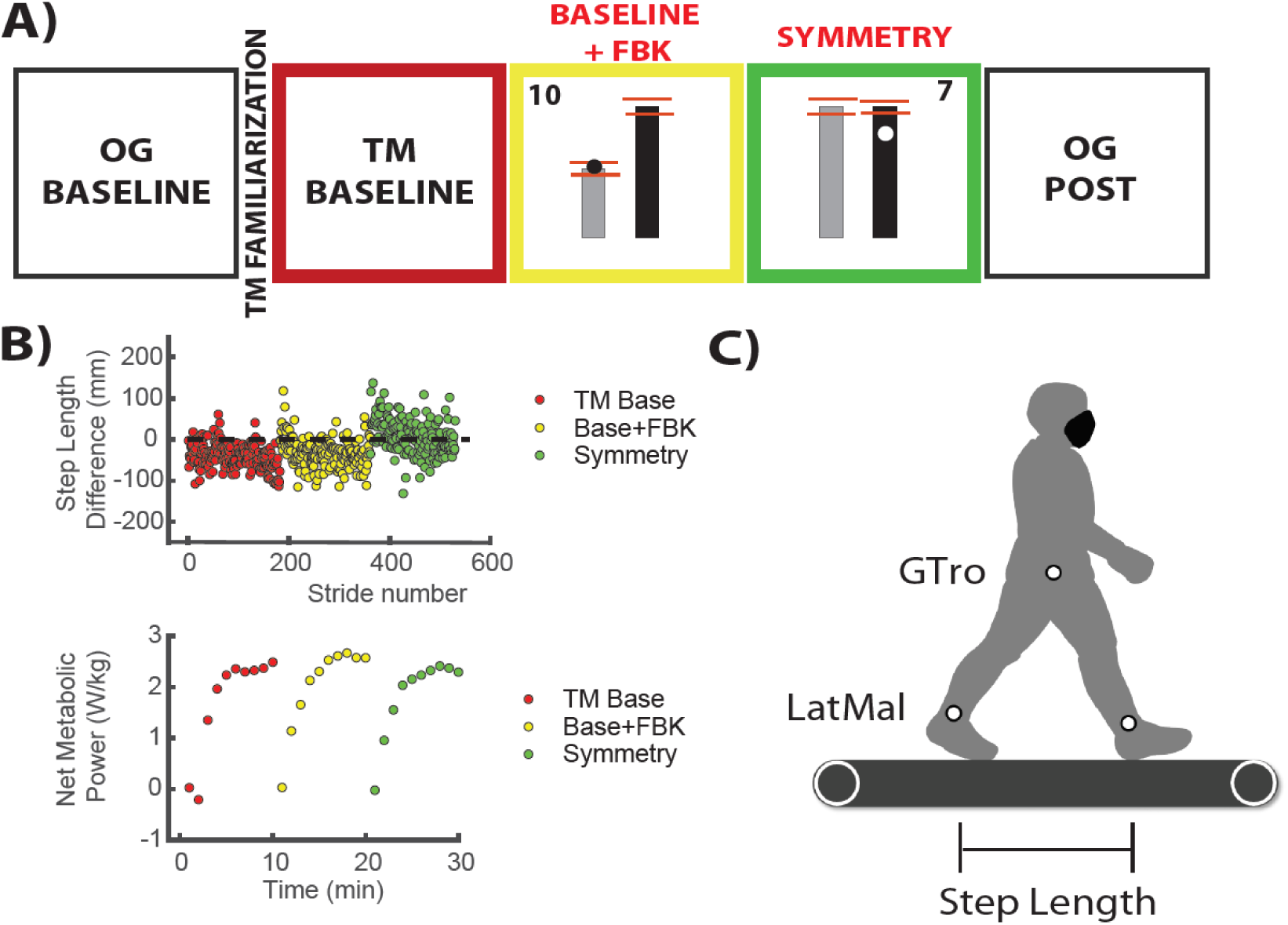
Experimental protocol and experimental setup. A) Experimental protocol and example of visual feedback. OG: Overground. TM: Treadmill. The red lines indicated the target window for the visual feedback trials which was defined as the twice the standard deviation of the step length difference measured during baseline. An example of the score is provided on the top left and right corners of the window. This score was a function of the distance of the marker to the target at foot strike. The score was updated on each side for every step. B) Raw step length difference data and metabolic power during TM trials. Data are color matched to the trials in the experimental protocol. Metabolic data were measured only for TM trials during *BASELINE, BASELINE+FBK, and SYMMETRY*. These trials lasted five minutes to ensure a steady metabolic cost. C) Experimental setup and marker locations.

Participants were provided with visual feedback of their step lengths via a display that was controlled by custom software written in Vizard (Worldviz, Santa Barbara, CA). The visual feedback consisted of two vertical bars specifying the desired length of their right and left steps. The real-time location of markers placed on their ankles was projected onto the vertical bars. Participants were instructed to walk such that the position of the ankle marker coincided with the top of the bars on the corresponding side at foot strike. Points were awarded to participants as follows (rounded to the nearest integer):

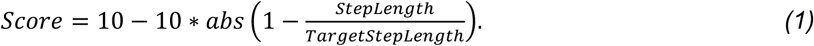

This formula was used to provide a score that varied systematically with the error between the desired and actual step length. Participants were verbally encouraged to obtain 10 points for every step on each side.

### Data Acquisition

#### Kinematic data

Kinematic data for passive markers placed bilaterally on the lateral malleoli and greater trochanters were collected using a 10 camera Qualisys Oqus system (QTM, Sweden). Foot strike and lift-off were estimated from peak anterior and posterior lateral malleoli excursions, respectively^28^.

#### Metabolic cost

Metabolic cost was assessed using expired gas analysis. Expired gas was sampled on a breath-by-breath basis, and the rate of O_2_ and CO_2_production were measured using a TrueOne® 2400 system (Parvomedics, UT). Substrate utilization during the experiment was determined using the Respiratory Exchange Ratio (RER), which is the ratio of carbon dioxide produced to the oxygen consumed. Metabolic power was assessed using a standard equation^29^. The average metabolic power from a standing baseline trial collected before the waking trials was subtracted from measurements made during all subsequent walking periods to yield net metabolic power. Metabolic data were supplemented with self-reported Ratings of Perceived Exertion (RPE)^30^, which were collected after each trial in 18 of 24 participants.

### Data Processing and Analysis

#### Step length asymmetry

Individual step lengths were defined as the fore-aft distance between the lateral malleoli at the time of the respective limb’s footstrike^7,25,31^. We characterized step length asymmetry using the magnitude of the difference in step lengths (|*SL*_*Diff*_*|*), defined as the non-paretic step length minus the paretic step length, and the direction of this difference, such that -1 indicated a longer paretic step and 1 indicates a shorter paretic step. Average values of *SL*_*Diff*_ were obtained for the last two minutes of walking from the five-minute trials.

#### Spatial and temporal contributions to step length asymmetry

We also expressed *SL*_*Diff*_ as the sum of spatial and temporal contributions to step length asymmetry, as previous work has shown that the variance in the metabolic cost of walking post-stroke can be partially explained by differences in paretic and non-paretic foot placement relative to the body (step position contribution, Equation 3)^8^:

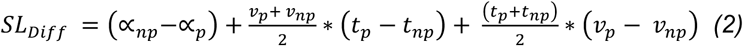

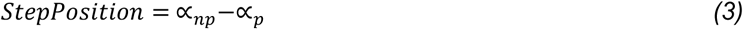

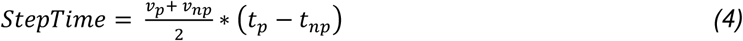

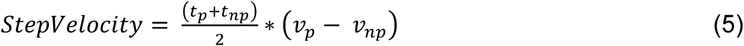

Here, α_*np*_ is a spatial variable indicating the difference in how far the non-paretic foot is placed in front of the body relative to the previous paretic foot placement, and α_*p*_ indicates the difference in how far the paretic foot is placed in front of the body relative to the previous non-paretic foot placement. v_*np*_ and v_*p*_ are the speed of the body relative to the non-paretic and paretic foot during stance. t_*np*_ and t_*p*_ are the non-paretic and paretic step times where t_*np*_ it is the time from non-paretic foot strike to paretic foot strike and vice versa for t_*p.*_ A detailed derivation of these equations can be found elsewhere^7^. Average values for each variable were calculated for the last two minutes in each trial.

#### Cost of Transport

The net metabolic power and RER corresponding to *BASELINE, BASELINE+FBK* and *SYMMETRY* were calculated from the averages during the last two steady-state minutes of each trial, consistent with the period over which spatiotemporal variables were analyzed. All measures of metabolic power were normalized by body mass and treadmill speed to obtain the net metabolic cost of transport (CoT), expressed in J/kg^*^m.

#### Statistical analyses

All statistical analyses were performed in Matlab R2016b (Mathworks, Natick, MA, USA). Data were tested for normality using the Kolmogorov-Smirnov test. If the data satisfied the normality test, repeated measures analyses of variance (RM-ANOVA) were implemented to test whether the values of the following variables differed across trials: |*SL*_*Diff*_*|*, step lengths, stance duration, swing duration, stride length, cadence, position, time and velocity contributions to *SL*_*Diff*_, CoT, RER, and RPE. The direction of the asymmetry was defined as a between-subjects effect in the repeated measures analyses. If data did not satisfy the normality requirement, repeated measures analyses were implemented using either the Wilcoxon sign rank test for paired data or the Friedman test. We verified that data satisfied the sphericity assumption using the Mauchly test. If the sphericity assumption was not satisfied, we used the Huyn-Feldt corrected p-value. Post-hoc comparisons were performed using the Tukey-Kramer correction for multiple comparisons. The significance level was set at p=0.05.

#### Regression analyses

We used robust regression to fit linear models relating participant-specific modifications in asymmetry or modifications in metabolic cost and spatiotemporal variables characterizing each participant’s walking pattern. We explored whether the dependent variables, change in asymmetry magnitude (Δ*SL*_*Diff*_) and change in CoT (ΔCoT) during *SYMMETRY* relative to *BASELINE+FBK* were associated with the following independent variables: magnitude of baseline *SL*_*diff*_, step position, step time, and step velocity contributions to *SL*_*Diff*_, and baseline asymmetry direction (*Asym*_*Positive*_). *Asym*_*Positive*_ is a binary variable that takes a value of 1 if the asymmetry is positive (shorter paretic steps) and 0 if the asymmetry is negative (longer paretic steps).

For each regression analysis, we first calculated variance inflation factors (VIF) to determine the candidate variables considered for inclusion in the model. If the VIF for a given variable was greater than 4, the variable was removed from the model to avoid multicollinearity. Once the set of candidate variables for each model were determined, all possible models with combinations of an intercept, linear combinations of the dependent variables, and two-way interactions between the independent variables were fit in Matlab using robust linear regression. The model with the lowest Akaike Information Criterion^32^ (AIC) was then selected as the best-fit model, and the quality of the model’s fit to the data was determined using a leave-one-out cross-validation (LOOCV) approach for computing R^2^.

## Results

### Effects of augmented visual feedback on spatiotemporal variables and CoT

We first determined whether the provision of visual feedback during *BASELINE+FBK* had an effect on any spatiotemporal variables relative to *BASELINE*. We observed a significant main effect of trial type on *|SL*_*Diff*_*|* (RM-ANOVA, F_2,44_=10.44, p<0.001, Figure 2A), but there was no difference in *|SL*_*Diff*_*|* during *BASELINE+FBK* compared to *BASELINE* (p=0.437). There were also significant main effects of trial type on paretic (RM-ANOVA, F_2,44_=6.141, p=0.009) and nonparetic step lengths (RM-ANOVA, F_2,44_=22.967, p<0.001). There were no differences in paretic step length during *BASELINE+FBK* compared to the *BASELINE* (p=0.335), but the non-paretic step length was 16 ± 10mm longer during *BASELINE+FBK* (p=0.033). The remaining spatiotemporal variables (stride length, cadence, paretic swing time, and non-paretic swing duration) did not differ during *BASELINE+FBK* compared to *BASELINE*, with the exception of stance time. Paretic and non-paretic stance were both 0.06s ± 0.02s longer during *BASELINE+FBK* compared to *BASELINE* (p=0.026 and p=0.025 respectively). Thus, other than slight increases in non-paretic step length and in paretic and non-paretic stance time, there were no apparent changes in spatiotemporal coordination induced by the precise foot placement requirements associated with the visual feedback.

**Figure 2.**
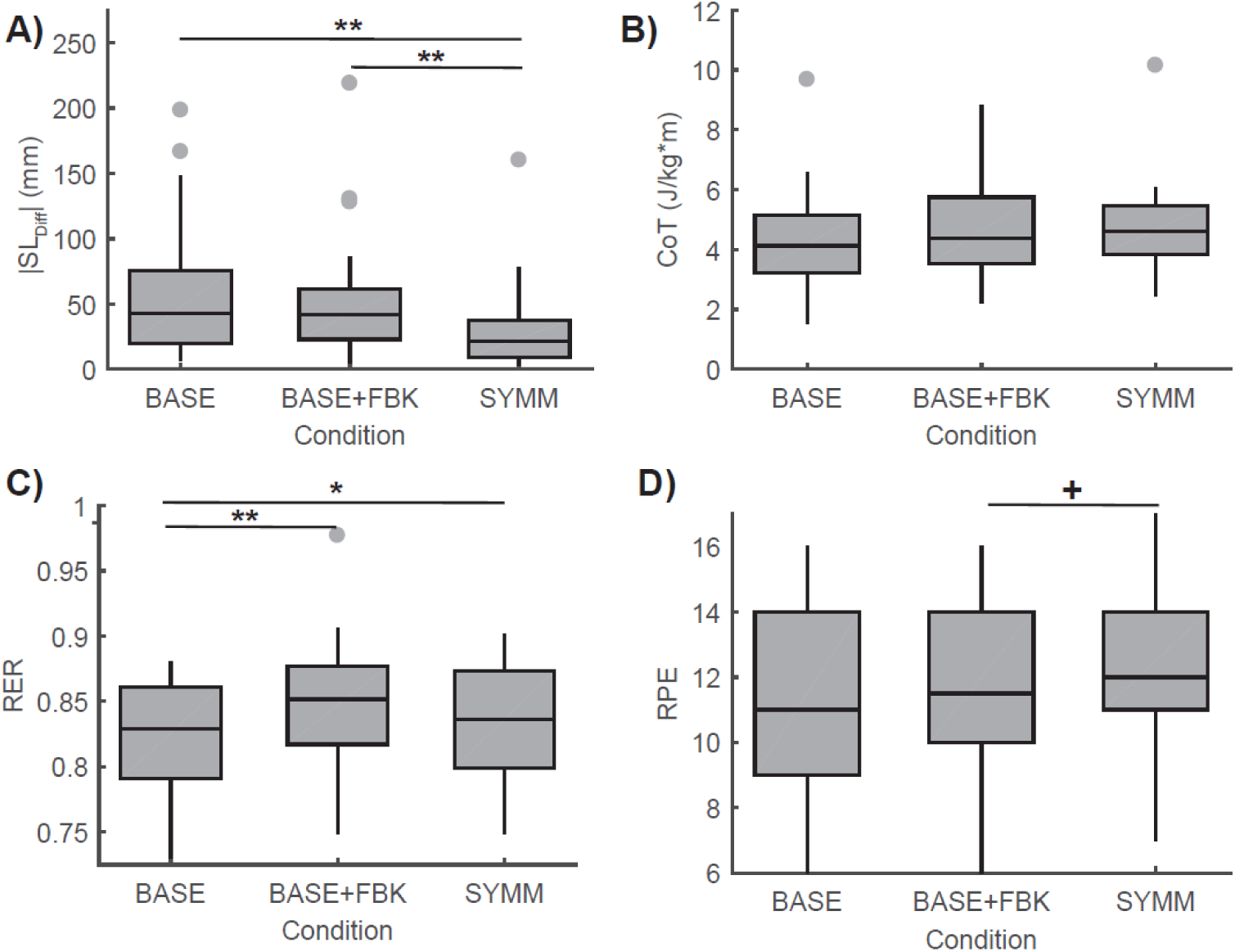
Effects of visual feedback on spatiotemporal control and metrics of energetic cost. A) Step length difference (SL_diff_) magnitude. There were no significant differences in step length difference between *BASELINE* and *BASELINE+FBK*. The feedback during *SYMMETRY* led to significantly smaller asymmetries compared to *BASELINE* and *BASELINE+FBK*. B) Cost of transport (CoT). No significant differences in CoT were observed across trial types. C) The respiratory exchange ratio (RER) increased due to the added feedback but did not increase further during the SYMMETRY condition. D) The rating of perceived exertion (RPE) increased marginally (F2,32=2.892, p=0.084, post-hoc ^+^p=0.077) when participants were instructed to generate symmetric step lengths. ^+^p<0.1, *: p<0.05, **:p<0.005.

Visual feedback of the desired and actual step lengths had contrasting effects on estimates of energetic cost. No significant differences in CoT were observed across conditions (RM-ANOVA, F_2,44_=2.293, p=0.123, Figure 2B). This is contrary to previous results in healthy, young participants where added feedback increased the metabolic cost by 18%^33^. The RER did vary across conditions (RM-ANOVA, F_2,44_=11.319, p<0.001, Figure 2C) and was higher during *BASELINE+FBK* relative to *BASELINE* (p=0.001), indicating that the added precise foot placement requirements were associated with increased CO_2_ production. Finally, there was a marginally significant effect of condition on the RPE (RM-ANOVA, F_2,32_=2.829, p=0.084, Figure 2D).

### Effects of symmetry training on spatiotemporal variables and CoT

When participants were provided with visual targets to lengthen their shorter step, there was a systematic reduction in |*SL*_*Diff*_*|* and simultaneous changes in other spatiotemporal variables. Average |*SL*_*Diff*_*|* during *SYMMETRY* was 30 ± 32mm compared to 54 ± 48mm during *BASELINE+FBK* (p=0.004, Figure 2A). There were no systematic effects of condition on the step position (RM-ANOVA, F_2,44_=2.546, p=0.104), step time (RM-ANOVA, F_2,44_=1.454, p=0.244), or step velocity (RM-ANOVA, F_2,44_=0.271, p=0.724) contributions to *SL*_*Diff*_. This indicates that participants used multiple, idiosyncratic strategies to reduce asymmetry.

There were no systematic changes in energetic cost during the *SYMMETRY* condition relative to *BASELINE+FBK*. The reduction in asymmetry did not lead to significant changes in the CoT (p=0.760, Figure 2B) or RER (p=0.182, Figure 2C). Although there was a trend toward participants having a higher perception of effort during *SYMMETRY*, this did not reach statistical significance (p=0.077, Figure 2D).

### Behavioral variables associated with the capacity to reduce asymmetry voluntarily

Since the direction and magnitude of *SL*_*Diff*_ were not homogenous across our sample (Figure 3A), we next explored how these variables influenced participants’ ability to reduce |*SL*_*Diff*_*|*. No differences in Fugl-Meyer scores (p=0.613), walking speed (p=0.394) or stride length (p=0.227) were observed between these groups. We found that the capacity to modify asymmetry depended on the magnitude of baseline *SL*_*Diff*_, the position and velocity contributions to *SL*_*Diff*_, and the interaction between position and velocity contributions and asymmetry direction (Table 2, Figure 3B-D). The model had an R^2^ of 0.68. For participants who took shorter paretic steps (*Asym*_*Positive*_=1), the net effects of the step position and velocity contributions were negligible, indicating that their ability to reduce asymmetry is mostly proportional to their baseline asymmetry. In contrast, for participants in the longer paretic group (*Asym*_*Positive*_=0), having a greater position contribution to *SL*_*diff*_ was associated with greater reductions in asymmetry while the velocity contribution reduces the capacity to reduce asymmetry. For 8/10 participants in the longer paretic group, the net effect of the position and velocity contributions was positive, resulting in additional reductions in Δ*SL*_*Diff*_ for this group.

**Figure 3.**
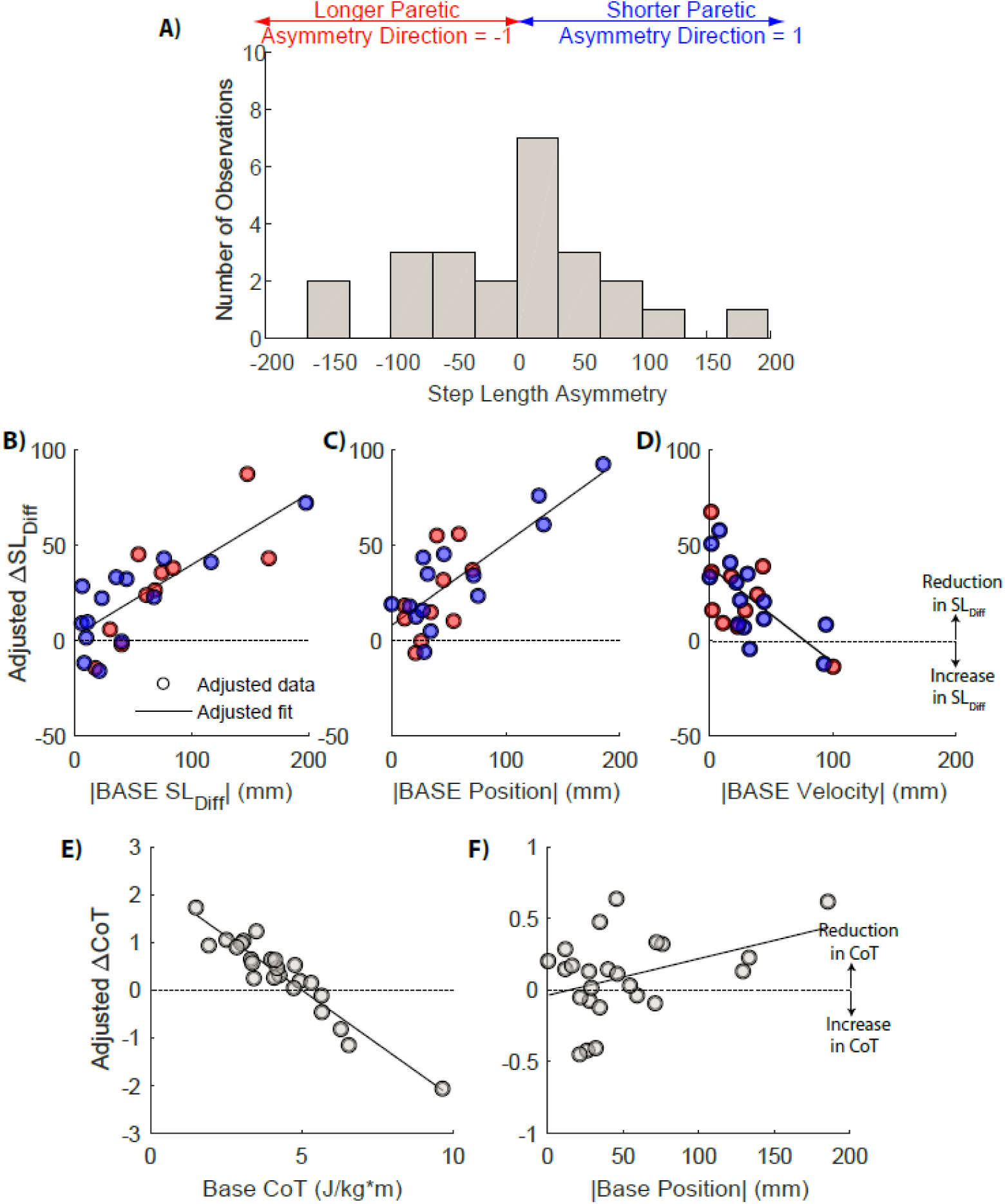
Baseline step length asymmetry and changes in asymmetry and metabolic cost. A) Histogram of the distribution of step length asymmetries for our sample of N=24 participants. B-D) Adjusted response plots for the main effects in the model describing associations between the change in SL_Diff_ and baseline asymmetry magnitude, position and velocity contributions (Table 3). A positive ΔSLDiff indicates a decrease in asymmetry. Participants in the longer paretic group are colored in red, whereas participants in the shorter paretic group are colored in blue. E-F) Adjusted response plots for significant main effects in the model in Table 4. A decrease in CoT during *SYMMETRY* corresponds to a positive ΔCoT.

**Table 2:**
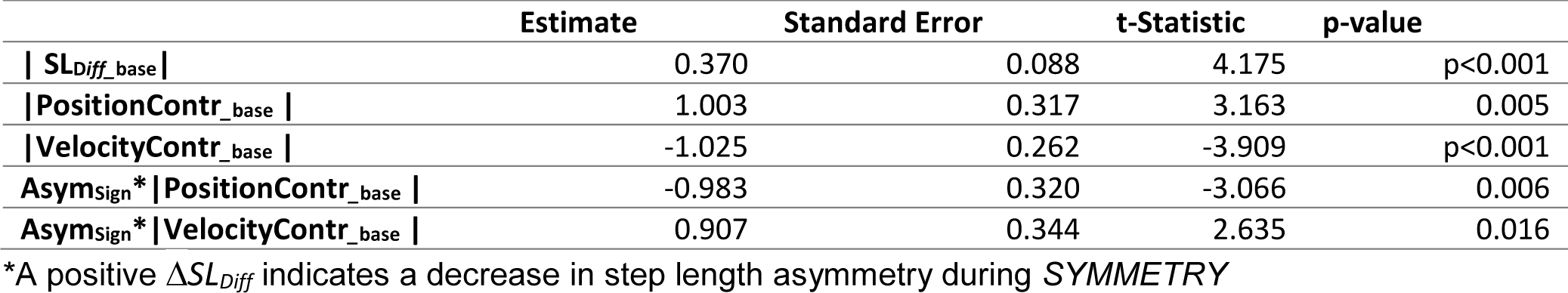
Parameter values for model relating Δ*SL_Diff_*\* to baseline spatiotemporal variables

We also found that changes in *SL*_*Diff*_ and individual step lengths during the *SYMMETRY* condition were associated with the direction of baseline asymmetry. For the longer paretic group, asymmetry was reduced by 36 ± 11mm during *SYMMETRY* (p=0.007). This was achieved by an increase in non-paretic step lengths of 53 ± 9mm compared to *BASELINE+FBK* (p<0.001). In contrast, for the shorter paretic group, asymmetry was not significantly different from *BASELINE+FBK* (average reduction of 14 ± 9mm, p=0.259) despite an increase in their paretic step length by 19 ± 5 mm (p=0.012).

### Associations between baseline spatiotemporal variables and changes in the CoT during SYMMETRY

Although we did not observe a significant main effect of condition on CoT, we found that the change in CoT from *BASELINE+FBK* to *SYMMETRY* was explained by individual differences in multiple baseline spatiotemporal variables (Table 3, Figure 3E-F). The linear model had an R^2^ of 0.72. The baseline CoT, which contributes to multiple terms of the model, had a negative net effect indicating that participants whose walking was already costly were likely to increase CoT (Figure 3E). This likely reflects the effects of impairment as there was a significant correlation between CoT and Fugl-Meyer (r = -0.52). Also, participants who had a larger position contribution to *SL*_*diff*_ were more likely to reduce CoT during *SYMMETRY* (Figure 3F).

**Table 3:**
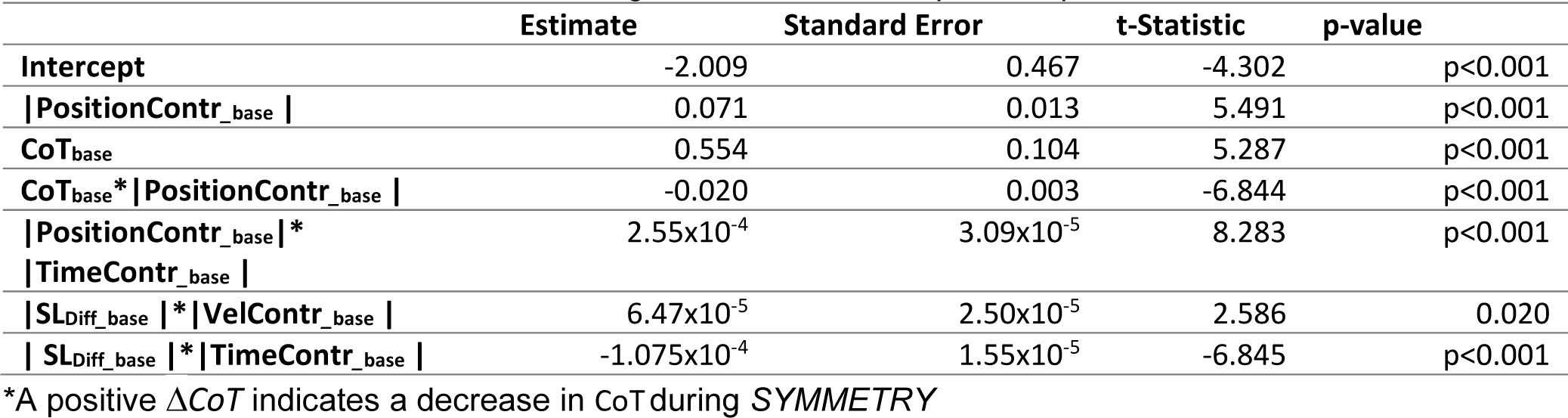
Parameter values for model relating Δ*CoT*\* to baseline spatiotemporal variables

### Transfer of reductions in asymmetry to over-ground walking

Out of 24 participants, 16 remained less asymmetric during *OG POST* compared to *OG BASE*. At the group level, there were no significant differences in *SL*_*diff*_ magnitude during *OG POST* compared to *OG BASELINE* (paired t-test, p=0.753), but the change in *SL*_*Diff*_ during *OG POST* varied systematically with the magnitude of *OG BASELINE SL*_*Diff*_. A simple linear model showed that the reduction in *SL*_*diff*_ during OG POST was 25 ± 9 % of the *SL*_*Diff*_ measured during *OG BASELINE* (p=0.012). However, the R^2^ of the model was just 0.10. We also observed systematic changes in paretic and non-paretic step lengths during *OG POST* relative to *OG BASELINE*. Overall, there was a significant increase in paretic step length of 16 ± 31mm RMANOVA, p=0.018) for all participants. Participants in the longer paretic group also had a significant increase in non-paretic step length, which was the focus of the training, during *OG POST* 38 ± 45mm (RM-ANOVA, p=0.005). No changes in stance time during *OG POST* were observed for the non-paretic (Wilcoxon signed rank test, p=0.775) or the paretic extremity (paired-samples t-test, p=0.906).

## Discussion

We sought to determine the factors that predict individual differences in the capacity of people post-stroke to voluntarily reduce step length asymmetry and transfer these reductions to overground walking. We also assessed how changes in asymmetry impact the metabolic cost of walking. Our results demonstrate that when provided with the proper visual feedback, individuals post-stroke retain the capacity to reduce step length asymmetry voluntarily.

However, this capacity varied systematically with the direction of baseline asymmetry. Participants who took longer paretic steps had a greater capacity to reduce asymmetry than participants who took shorter paretic steps. Transfer of training to over-ground walking was also associated with the baseline direction of the asymmetry, with participants in the longer paretic group maintaining increases in the paretic and non-paretic step lengths, while participants in the shorter paretic group only maintained increases in paretic step length. Lastly, we found that reductions in CoT were negatively correlated with each participants’ baseline CoT such that more impaired participants (higher CoT) were likely to increase CoT in the SYMMETRY condition.

### Asymmetry direction influences that capacity to reduce step length asymmetry voluntarily

We found that a relatively complex model was necessary to explain the extent to which baseline features of each participant’s walking pattern could predict their ability to reduce asymmetry. Participants’ ability to reduce asymmetry voluntarily was positively correlated with their baseline *SL*_*Diff*_, as more asymmetric participants have a greater range for asymmetry reduction. Moreover, the effects of baseline step position and step velocity contributions differed based on participants’ baseline asymmetry direction. For participants in the longer paretic group, the step position and step velocity contributions to asymmetry were, respectively, positively and negatively correlated with the ability to reduce asymmetry. This may result from participants with large step velocity asymmetries having difficulty progress over the paretic limb, thus limiting their ability to reduce asymmetry. Group level analyses indicated that participants who took shorter paretic steps did not experience significant reductions in asymmetry. This may have resulted from deficits in paretic hip and knee flexion^5,13,15,34^ which would limit the capacity to lengthen paretic steps. This could also potentially be explained by the fact that over half of the participants in this group had negligible asymmetries. However, this was addressed in the regression model through the inclusion of a term representing baseline asymmetry magnitude. This suggests that the effects of asymmetry direction are independent of asymmetry magnitude.

### Individual differences in the effects of asymmetry reductions on metabolic cost

Although we did not observe consistent group-level changes in metabolic cost, we found that the both magnitude and direction of changes in metabolic cost depended on participants’ baseline CoT. Specifically, reductions in CoT occurred in those individuals who had a low baseline CoT and increases were observed in individuals who were most costly. This is likely explained by the fact that participants who had the highest CoT were most impaired and would, therefore, have more difficulty reducing asymmetry. In contrast, individuals who were less impaired may have achieved reductions in energetic cost that may have been due to improvements in symmetry. This suggests that the potential benefits of reducing asymmetry are not uniform across all stroke survivors. Attempts to reduce asymmetry in patients who are highly impaired may not be worth the added effort whereas less impaired individuals may benefit from approaches that focus on explicitly reducing step length asymmetry.

The lack of group-level reductions in CoT resulting from reductions in step length asymmetry was not consistent with previous studies which have shown that reductions in asymmetry resulting from 12 weeks of treadmill training with functional electrical stimulation (FastFES) are associated with reductions in metabolic cost ^34,36^. One potential explanation for differences between our results and those reported previously is that our participants had reductions in asymmetry that were ~4 times greater than those due to the FastFES intervention^34,36^. Therefore, it is possible that the associations reported previously may not hold for larger reductions in asymmetry. A second potential explanation is that reductions in metabolic cost following FastFES training were due to training-dependent increases in the functional capacity of the plantarflexors^5,34–36^. Whether similar effects would be observed in a longitudinal version of our study remains to be seen.

### Transfer of training to over-ground walking

Previous studies have examined the transfer of training from a treadmill to over-ground walking using resistive forces^37^ or using split-belt training^38^. In the study using resistive forces, participants transferred the increase in short step length to over-ground walking, but this was accompanied by a decrease in the long steps. This is contrary to our results as we saw an increase in the length of both steps for our longer paretic group and an increase in paretic step lengths for our shorter paretic group. Our results are more consistent with the transfer observed after split-belt training, with participants also showing an approximately 25% reduction in step length asymmetry after four weeks of training with 6/11 (55%) individuals becoming more symmetric after training compared with 16/24 (67%) in the current study.

One of the main limitations of our protocol is the use of a single session approach which may not have allowed for participants to become fully familiarized with the novelty of the task. However, we attempted to address this issue by providing a short practice period at the beginning of the session. In addition, by having only a single session of training, we could not capture the consolidation of the reductions in asymmetry that could have occurred during multiple sessions^40^. Another limitation is the fact that modifying a single gait variable inherently changes other variables. In our case, increasing the shorter step to match the long step length increases the overall stride length and reduces cadence. It is possible that the increase in metabolic cost due to the decreased cadence^39^ could have masked any potential reductions in metabolic cost during *SYMMETRY*.

In conclusion, we have shown that patient-specific differences in the capacity to voluntary reduce step length asymmetry can be largely explained by differences in baseline characteristics of their walking pattern. These differences likely stem from the heterogeneity of deficits in support, propulsion, and limb advancement on the paretic limb. The energetic effects of reductions in asymmetry also depended on the patient-specific impairment such that patients who are more impaired were likely to increase energetic cost when reducing step length asymmetry. Thus, our results highlight the need to characterize and account for inter-individual differences in locomotor function when considering the use of standardized training approaches^41,42^.

## Acknowledgments

We would like to thank all participants for volunteering for this study. We would also like to acknowledge Lindsey Trejo, Chang Liu and Aram Kim for assistance with data collection. Finally, we would like to thank Dr. Carolee Winstein, Dr. Sara Mulroy and Dr. Nicolas Schweighofer for their valuable feedback. This study was funded by the American Heart Association post-doctoral fellowship 16POST29610000 to N. Sanchez.

